# Cool ocean temperatures fail to buffer the impacts of heat exposure during low tide on the behaviour and physiology of a keystone predator

**DOI:** 10.1101/2024.03.07.584009

**Authors:** Lydia N. Walton, Viola R. Watts, Jasmin M. Schuster, Amanda E. Bates

## Abstract

Air temperatures are warming at faster rates than ocean temperatures, and this “land-sea warming contrast” may create reprieves from thermal stress by providing cool underwater refugia during extreme heat events. Here we tested the impacts of the “land-sea warming contrast” on physiology (metabolism) and behaviour (feeding) in the juvenile life stage of a keystone intertidal predator, *Pisaster ochraceus*, by experimentally manipulating air (∼20℃, 25℃, 30℃) and water (∼15℃, 20℃) temperatures (at independent rates) representing early summer, late summer, and heatwave conditions in Barkley Sound (British Columbia, Canada). We further made observations of air temperatures, sea surface temperatures, and *Pisaster* moribundity at our study location to support interpretation of our results. We predicted metabolism and feeding would increase with early and late summer temperatures, but decrease during heatwave conditions as animals surpass their thermal optimum. We observed the greatest mortality and lowest feeding in juvenile *Pisaster* exposed to cool ocean temperatures (∼15℃) and high aerial temperatures typical of extreme heat events (∼30℃). Feeding rates increased with heat stress duration, indicating animals may be compensating for elevated metabolism. Metabolic rates did not differ between air temperatures, but oxygen consumption was higher in animals with access to mussels than for *Pisaster* that were fasted. The highest levels of experimental and field moribundity were observed in August, indicating *Pisaster* may have accumulated physiological stress damage following elevated air and ocean temperatures throughout the summer. Our research implicates shifts in community dynamics due to the loss of this keystone species as air temperatures warm.

**Summary Statement:** Cooler ocean temperatures, rather than creating thermal refugia, may cause physiological stress for juvenile *Pisaster ochraceus* exposed to warm air during low tide.

## Introduction

Global surface temperatures have risen by ∼1°C in the past century, however, the rate of warming differs between terrestrial and marine systems (Byrne and O’Gorman, 2018; IPCC, 2023). Indeed, land surface temperatures (LST) are increasing at a faster rate than sea surface temperatures (SST) based on observations and climate models (Sutton *et al*., 2007; Joshi *et al*., 2008, 2013). This decoupling of sea surface and land surface warming patterns is distinguished as the “land-sea warming contrast” and has important implications for ecological communities, in particular those that use habitats in both terrestrial and ocean systems (Manabe *et al*., 1991).

Due to differences in warming rates and temperature variability between land and sea, many intertidal areas (*i.e.*, where the ocean and land intersect between high and low tides) already exhibit extreme temperature contrasts, with resident organisms experiencing temperature changes upwards of 20°C when alternating between submersion and emersion (Helmuth and Hofmann, 2001; McGaw *et al*., 2015). It is possible cooler ocean temperatures may provide thermal refugia from intense heat and solar radiation on land, and minimize lethal or sublethal heat stress as animals move down the shore (Dowd *et al*., 2015; Dong *et al*., 2017; Han *et al*., 2020). For instance, several species of mobile invertebrates occupy submerged or shaded microhabitats (*e.g.*, tide pools, under rocks) during low tide where body temperatures remain more stable and several degrees lower than in sun-exposed microhabitats (Helmuth and Hofmann, 2001; Szathmary *et al*., 2009; Sun *et al*., 2023). Another possibility is that the widening temperature difference between land and sea could increase the likelihood of species exceeding their thermal tolerance limits (*e.g.*, upper limit/heat tolerance). In response to seasonal temperature change or extreme climatic events (*e.g.*, heatwaves), organisms may shift their upper and lower tolerance limits to better accommodate ambient temperatures (Pörtner *et al*., 2006).

Here, we use the rocky shore intertidal of the Pacific Northwest as a model system for investigating the potential impacts of the “land-sea warming contrast” on ecological communities. The physiological and behavioural responses of key intertidal organisms (*e.g.*, ecosystem engineers, keystone predators) to different temperature exposures have been researched extensively and provide an important foundation for exploring the effects of novel climatic conditions on community dynamics (Sagarin *et al*., 1999; Harley *et al*., 2006; Monaco *et al*., 2016; Dong *et al*., 2017; Morón Lugo *et al*., 2020; Robles *et al*., 2021). More specifically, metabolic rates (*i.e.*, the rate at which organisms’ uptake energetic resources and allocate these to survival, growth and reproduction) vary predictably with temperature (Boltzmann, 1872) in ectotherms, increasing from a critical lower thermal limit (*CTmin*) up to a thermal optimum (*Topt*) before dropping towards a critical upper thermal limit (*CT_max_*) (Gillooly *et al*., 2001; Brown *et al*., 2004; Dell *et al*., 2011; Rebolledo *et al*., 2021). Therefore, a decreasing metabolic rate can indicate temperatures have surpassed *T_opt_* and are approaching critical thermal limits, a useful tool for predicting the impacts of extreme climatic events (*e.g.*, heatwaves) on ecological communities. Lowered metabolic rates in response to climate warming can also affect behaviour, with some organisms displaying reduced feeding under chronic heat stress so that more oxygen is available to processes other than digestion (Dahlhoff *et al*., 2001; Pincebourde *et al*., 2008).

Decreases in feeding rate can in turn affect community composition and species diversity if predators exhibit top-down control of an ecosystem (Paine, 1966). Thus, alterations in physiological and behavioural processes due to different exposures to warm ocean and air temperatures have important implications for community dynamics and ecosystem functioning, especially in intertidal systems.

We investigated the feeding (*i.e.*, rate of mussel consumption) and metabolic (*i.e.*, oxygen consumption) responses of an intertidal predator (ochre sea star; *Pisaster ochraceus*, hereafter *Pisaster*) to experimentally manipulated seawater and air temperatures representing early summer, late summer and heatwave conditions in Barkley Sound (British Columbia, Canada).

First, we examined changes in feeding rate when juvenile *Pisaster* were exposed to contrasting water and air temperatures in a simulated tidal cycle. We predicted that aerial heat stress would have a negative impact on mussel (*Mytilus* spp.) consumption, with suppressed feeding occurring at higher air temperatures near the upper thermal tolerance limits of *Pisaster* (Pincebourde *et al*., 2008). We also predicted that warmer water temperatures would help animals compensate for the energetic demands of aerial heat stress by increasing *Pisaster* metabolic rate, supporting greater feeding activity (Sanford, 1999; Fly *et al*., 2012). After observing feeding during the experiment, we further tested for a shift in feeding rate that related to the experimental time sequence, and hypothesized that rates of mussel consumption would increase significantly once heat stress was terminated. Next, we focused on the relationship between food availability, *Pisaster* metabolism, and air temperature. We predicted fasted organisms would suppress their metabolic rate, which is a sign of poor physiological condition and an indicator of increased susceptibility to thermal stress (Dahlhoff *et al*., 2001). As fasted organisms were expected to be more susceptible to thermal stress, we also predicted high levels of mortality in fasted *Pisaster* experiencing warmer air temperatures (∼30℃). After concluding both experiments, we compared mortality and feeding rates between treatments that were the same across the two experiments (*i.e.*, having similar food, seawater and air temperature conditions) to determine if differences existed between animals collected in early summer versus late summer. Finally, we made observations of air temperatures, sea surface temperatures, and *Pisaster* moribundity (*i.e.*, individuals in a state of dying) at our study location in early (May, June) and late (August) summer to determine average environmental conditions and to validate our experimental treatments. We predicted a higher prevalence of moribund *Pisaster* in late compared to early summer as small increases in temperature can result in greater spread and intensity of disease in ochre sea stars (Bates *et al*., 2009).

## Materials and Methods

We conducted two separate experiments at different times, one with the goal of testing *Pisaster* feeding response to contrasting water and air temperatures (Experiment 1, June 2023) and the second testing *Pisaster* metabolic response to food availability and air temperature (Experiment 2, August 2023). A new cohort of juvenile *Pisaster* was collected for each experiment. Field observations of *Pisaster* moribundity were conducted once per month in May, June and August 2023.

### Air and sea surface temperatures

Air and sea surface temperatures (SST) in Barkley Sound (British Columbia, Canada) supported our selection of the water and air temperatures in our experiments. Air temperatures from Eagle Bay (48°50’01.9”N, 125°08’47.7”W; **Fig. 1**) were recorded using an EnvLogger temperature sensor (version T2.4), while SST for the area was recorded by the Amphitrite Point lightstation in Ucluelet (48°55’16.1”N, 125°32’28.1”W; https://open.canada.ca/data/en/dataset/ 719955f2-bf8e-44f7-bc26-6bd623e82884). The maximum air temperature recorded at Eagle Bay in 2022 (∼30℃, July) was used to set the warmest air treatment (**Table S1**). The ∼20℃ and ∼25℃ experimental air treatments were selected based on midday low tide conditions recorded at three sites (Eagle Bay, Grappler Narrows, Strawberry Point; **Fig. 1**) near Bamfield, British Columbia, Canada in early summer (*i.e.*, May, June) and late summer (*i.e.*, July, August) respectively (**Table 1**). The cooler water treatment (∼15℃) represented SST in early summer while the warmer water treatment (∼20℃) generally represented SST in late summer (**Tables S1, S2**).

**Fig 1.**
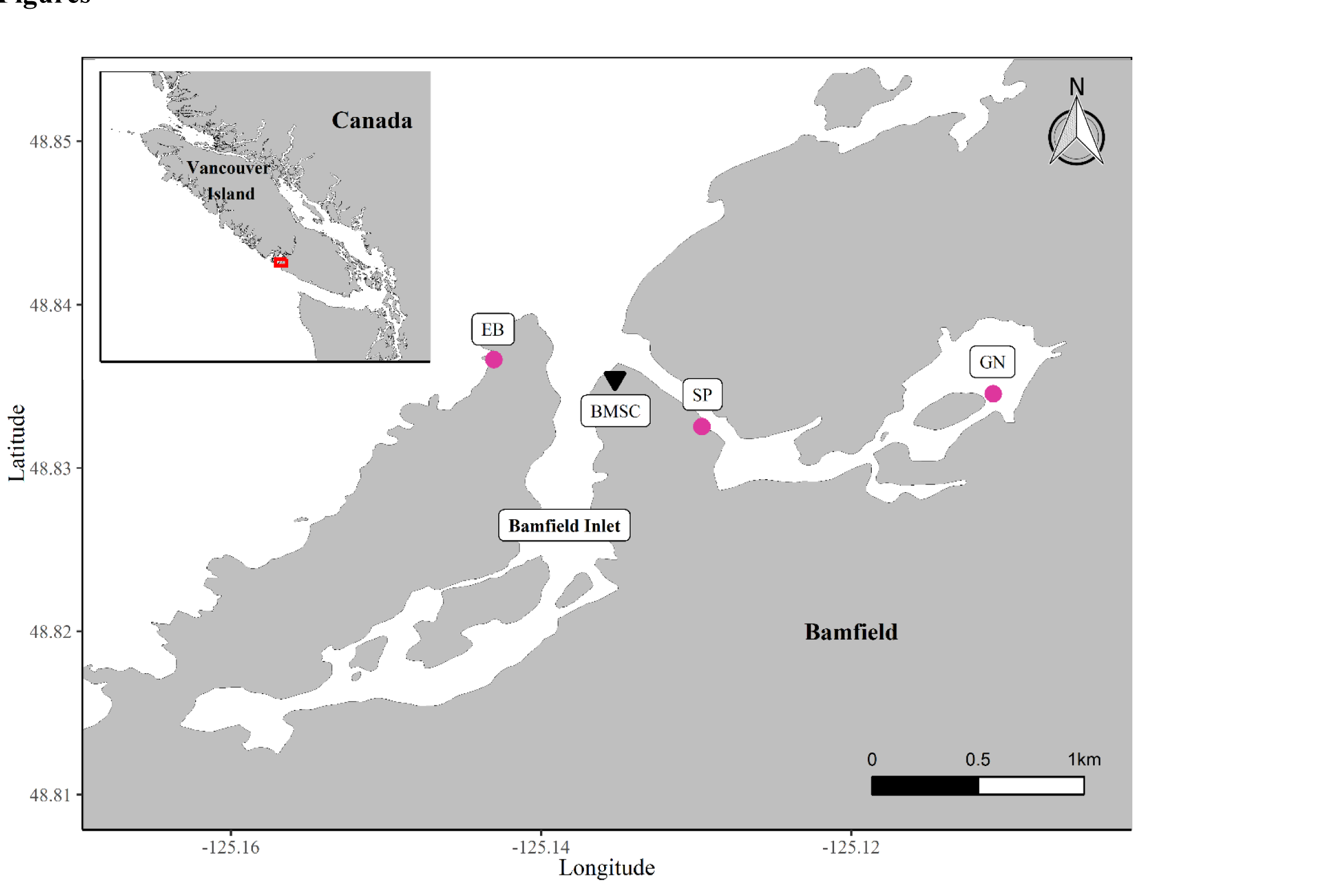
Map of three localities (pink circles) near the Bamfield Marine Sciences Centre (BMSC; black triangle) in British Columbia, Canada used for field observations in the summer of 2023. All three sites (Eagle Bay (EB), Strawberry Point (SP), and Grappler Narrows (GN)) were surveyed for the presence of moribund adult and juvenile *Pisaster ochraceus*. Eagle Bay was selected as the collection site for juvenile ochre stars used in experiments (June and August 2023). An EnvLogger temperature sensor was deployed at EB to record air temperatures in the intertidal zone (Baum Lab research group, University of Victoria).

**Table 1.** The percent of moribund *Pisaster ochraceus* observed at three sites (Eagle Bay, Grappler Narrows, and Strawberry Point) near Bamfield, British Columbia in May, June and August 2023.

| Site (n = sample size) | Month | Moribund adult (%) | Moribund juvenile (%) | Air temp (°C) | SST (°C) |
| --- | --- | --- | --- | --- | --- |
| **Eagle Bay (n = 247 adults, 5 juveniles)** | May | 0.8 | 0 | 15.4 | 14.7 |
| **Eagle Bay (n = 332 adults, 13 juveniles)** | June | 0.6 | 0 | 21.8 | 16.7 |
| **Eagle Bay (n = 198 adults, 48 juveniles)** | August | 9.6 | 8.3 | 21.1 | 19.5 |
| **Grappler Narrows (n = 38 adults, 5 juveniles)** | May | 0 | 0 | 17.8 | 15.2 |
| **Grappler Narrows (n = 56 adults, 61 juveniles)** | June | 0 | 0 | 19 | 14.6 |
| **Grappler Narrows (n = 79 adults, 70 juveniles)** | August | 5.1 | 2.9 | 23.5 | 17.6 |
| **Strawberry Point (n = 76 adults)** | May | 0 | 0 | 21 | 16.7 |
| **Strawberry Point (n = 50 adults, 1 juvenile)** | June | 0 | 0 | 14.3 | 14.7 |
| **Strawberry Point (n = 59 adults, 1 juvenile)** | August | 0 | 0 | 18.3 | 17.5 |

*Pisaster ochraceus* were counted along walking belt transects at each site during a midday low tide. The percentage of moribund adults and juveniles was recorded. Additionally, the air temperature and sea surface temperature (SST) at each site was recorded at the time of the survey (INKBIRD IBS-TH2 Plus temperature probe). SST was recorded at the low tide line.

### Experiment 1 (June). Effects of the “land-sea warming contrast” on Pisaster feeding rate

#### Collections

Juvenile *Pisaster ochraceus* (hereafter *Pisaster;* n = 72) were haphazardly hand collected from Eagle Bay, Bamfield, British Columbia (48°50’01.9”N, 125°08’47.7”W; **Fig. 1**) in early June 2023 and transported by boat (in water-filled buckets) to the Bamfield Marine Sciences Centre (BMSC; **Fig. 1**) where animals were housed in flow-through seawater tables at ambient temperature (∼13℃, **Table S2**). Specimens were measured upon collection, including arm length (measured from the centre of the aboral disc to the tip of the arm ray closest to the madreporite in mm), disc diameter (diameter of the aboral disc in mm), and wet mass (g). All individuals had a wet mass between 1 g and 30 g (mean ± SD = 8.2 ± 6.5 g) and were classified as juveniles based on size (*Pisaster* reach sexual maturity at wet masses of 70 to 150 g; Mauzey, 1966; Menge and Menge, 1974; Robles, 2013). Mussels (*Mytilus* spp.) were collected from docks and pilings within the Bamfield Inlet (**Fig. 1**) and had shell lengths ranging from 6.4 mm to 41.1 mm (mean ± SD = 21.3 ± 6.47 mm, n *=* 783) and shell widths ranging from 4.4 mm to 25.5 mm (mean ± SD = 11.9 ± 3.47 mm, n *=* 783).

#### Experimental set-up

Juvenile *Pisaster* were randomly assigned to one of six treatments (n = 12 per treatment) crossing different water (15 and 20°C) and air (20, 25, 30°C) temperature exposures (respectively, water/air temperatures in °C: 15/20, 15/25, 15/30, 20/20, 20/25, 25/30). The difference in mean body size (mean disc diameter) between each treatment was less than 3 mm.

A set of four sea tables was used for heat-stress experiments, with each sea table (volume 40 L) supplied continuously with flow-through seawater at a rate of ∼0.625 L min^-1^. Two sea tables were heated to ∼15°C using 200 W aquarium heaters (AEH3617090 Jager aquarium thermostat heater, Eheim) and the remaining two sea tables were heated to ∼20°C using 250 W aquarium heaters (3618090 Jager aquarium thermostat heater, Eheim). Animals were housed in individual aquaria (volume 750 mL) with 18 aquaria placed in each sea table and swapped systematically between sea tables daily, preserving treatment conditions to minimize the possibility of sea table effects (*i.e.*, individuals in the 15°C water treatment alternated between both 15°C water sea tables).

#### Experimental periods: adjustment, heat stress, and recovery

Prior to treatment start, animals were fully submerged in ambient seawater (∼13°C) for eight days to adjust to laboratory conditions. During this adjustment period (days 1-8) *Pisaster* were fasted to ensure all animals were in a similar physiological condition prior to heat stress. Heat stress was applied on day nine, and ochre stars were cycled between an 18-hour submersion in sea stables heated to ∼15°C or 20°C water temperature and a 6-hour aerial emersion in temperature-controlled incubators heated to ∼20°C, 25°C or 30°C air temperature for 16 consecutive days. Temperature probes (INKBIRD IBS-TH2 Plus) recorded the water temperature of each sea table for the duration of the experiment (32 days) and in each incubator for the duration of the heat stress period (16 days) to ensure treatment conditions were maintained (**Table S2**). *Pisaster* were fed mussels *ad libidum* (*i.e.*, minimum 3 mussels in every aquarium at any given time) throughout the heat stress period, and empty shells (*i.e.*, consumed mussels) were removed and counted every 24 hours. *Pisaster* survival was monitored and moribund individuals were removed from the aquaria every 24 hours. Following the heat stress period, sea table temperatures were returned to ambient (∼13 °C, **Table S2**) and animals remained fully submerged. This recovery period lasted for eight days with an *ad libidum* feeding regime and empty mussel shells were removed and counted daily. All surviving ochre stars were returned to the collection site at the end of the experiment.

#### Statistical analysis

Analyses of *Pisaster* survival and feeding activity were conducted in R version 4.1.2 (R Core Team, 2021). The survival probability of *Pisaster* and consumed *Mytilus* spp. was determined for each treatment combination using Kaplan-Meier survival curves and Cox proportional hazards models from the “survival” and “survminer” packages (Kassambara *et al*., 2021; Therneau *et al*., 2023). The effect of contrasting water and air temperatures (*i.e.*, water temperature and air temperature treatment combinations) on *Pisaster* feeding activity was assessed using generalized linear mixed models (“glmmTMB” package; Brooks *et al*., 2017) with Poisson distributions. The number of mussels consumed by *Pisaster* was compared between treatments across the entire experimental period (*i.e.*, heat stress and recovery period) as well as for each 8-day “time block” of the experiment (*i.e.*, first half of the heat stress period: days 1-8; second half of the heat stress period: days 9-16; recovery period: days 17-24). A change point analysis (“changepoint” package; Killick *et al*., 2022) determined if a shift in feeding activity occurred during the transition from the heat stress period to the recovery period by detecting differences in the mean number of mussels consumed by *Pisaster* for each treatment. Only sea stars that survived and retained five intact arms were included in the feeding activity analyses (n

= 63).

### Experiment 2 (August). Food availability, Pisaster metabolism, and air temperature

#### Collections and experimental design

Juvenile *Pisaster* were collected in early August 2023 (Eagle Bay, as above) using the same methods as the feeding activity experiment (Experiment 1). All ochre stars had a wet mass between 1 g and 18 g (mean ± SD = 6.39 ± 4.33 g, n = 56) and were classified as juveniles based on size (Mauzey, 1966; Menge and Menge, 1974; Robles, 2013). Animals were randomly assigned to one of four treatments (n = 14 per treatment) crossing different food (+ mussels and - mussels) and air (25°C and 30°C) temperature exposures (respectively, food/air temperatures: +mussels/25°C, +mussels/30°C, -mussels/25°C, -mussels/30°C). *Pisaster* in the +mussels treatments were provided mussels *ad libidum* throughout the experiment while individuals in the -mussels treatments were given no access to food (*i.e.*, fasted). Mussels (*Mytilus* spp.) were collected from docks and pilings within the Bamfield Inlet (**Fig. 1**) and had shell lengths ranging from 6.2 mm to 32.3 mm (mean ± SD = 15.0 ± 5.65 mm, n *=* 96) and shell widths ranging from 3.8 mm to 18.8 mm (mean ± SD = 8.6 ± 3.25 mm, n *=* 96). This experiment consisted of two parts: **(1)** a 16-day period during which individuals acclimated to food/air treatments and **(2)** an experimental estimation of metabolic rates for all surviving ochre stars using a closed respirometry system.

#### Acclimating to food and air treatments

A set of four sea tables and two temperature-controlled incubators (as described in Experiment 1) were used during the acclimation period of the experiment, with sea tables heated to 20°C and incubators heated to 25°C or 30°C. Temperature probes (INKBIRD IBS-TH2 Plus) remained in each sea table and incubator for the duration of the experiment with temperatures recorded daily (**Table S2**). Following collection, *Pisaster* were submerged in the heated sea tables for 24 h to recover from handling stress. Animals were then alternated between sea tables and incubators in a simulated tidal cycle (as described in Experiment 1) that was repeated for 16 consecutive days. Empty mussel shells were counted every 24 h to determine if there were any differences in feeding activity between the air treatments. To mitigate sea table effects, *Pisaster* aquaria were swapped systematically between sea tables corresponding to the same feeding treatment. Following the acclimation period, ochre stars remained submerged in heated sea tables until metabolic testing began.

#### Experimental assay system for measuring oxygen consumption

A custom-built experimental system was used to measure the oxygen consumption (ṀO_2_) of individual *Pisaster* after the acclimation period was completed (described in detail: Schuster *et al*., 2022; Schuster and Bates, 2023). In brief, the system consisted of a table with 10 removable acrylic chambers (240 mL, 6.5 cm in diameter) with magnetic stir plates, placed inside a temperature-insulated seawater tank. The tank was equipped with a heater, chiller and water pump that circulates temperature-controlled seawater across the tank, ensuring consistent temperatures across the 10 chambers. The seawater in the tank was continuously aerated by an air pump attached to an air stone, and each chamber was fitted with its own temperature and fiber-optic oxygen dipping probe (PreSens Pt1000 and DP-PSt7-10-L2.5-ST10-YOP). Measurements of temperature and oxygen level (% saturation) were made every second via PreSens software (PreSens Measurement Studio 2, Version 3.0.3; Precision Sensing GmbH, Regensburg, Germany), and oxygen measurements were automatically temperature-corrected by the software. Oxygen probes were calibrated prior to oxygen consumption measurements, with air saturated, room temperature seawater (100 % O_2_) and room temperature distilled water containing sodium sulfite (no O_2_; 1 g Na2SO3 dissolved in 100 mL of water).

#### Pre-measurement procedures

To estimate the routine metabolic rate at the end of the acclimation period, the ṀO_2_ of *Pisaster* was measured using closed respirometry with the system described above. A maximum of nine individuals could be assayed per day (due to space constraints of the respirometry system), thus measurements of ṀO_2_ for all surviving individuals (n = 18) were staggered over two assay days. On the evening before the assay, the mass and volume of each individual was measured by displacement [volumetric displacement = volume of seawater with ochre star - volume without ochre star]. Animals were randomly drawn from all treatments and distributed individually into each of the nine chambers, while the tenth chamber remained empty to account for background respiration (*i.e.*, due to microorganisms). The chambers were covered with mesh- fabric to prevent escape and placed into a holding tank overnight. The holding tank was supplied with filtered (10 µm) flow-through seawater at the treatment temperature (∼20°C). Oxygen consumption measurements were started the subsequent morning.

#### Oxygen consumption measurements

Oxygen consumption was measured for all ochre stars at the treatment water temperature (∼20°C). The insulated cooler was filled with fresh, filtered (10 µm) seawater and maintained at the treatment temperature for each run. To initiate a measurement run, the lids to each chamber (9 ochre stars, one blank) were sealed. Oxygen consumption (ṀO_2_ in % air saturation) was recorded until oxygen levels dropped by 5 to 10%. The total measurement time varied from 30 min to approximately 1.5 h, with longer measurement times for *Pisaster* with lower masses of metabolically active tissue. Once the measurements were completed for each of the 10 chambers, all ochre stars included in the measurement run were removed from their chambers and immediately frozen prior to ash-free dry mass (AFDM) determination. The chambers were emptied and cleaned. The same evening, nine fresh organisms were placed into the chambers (after having their mass and volume measured) and placed into the overnight holding tank, repeating the same measurement procedure as before. The seawater tank was emptied, rinsed with warm freshwater, and refilled with fresh seawater to prepare for the next assay day.

#### Ash-free dry mass (AFDM)

*Pisaster* AFDM was determined to quantify the amount of organic tissue, or metabolically active tissue, of each ochre star. Empty aluminum weigh boats were placed in a muffle furnace (500°C) for 12 h to remove any organic matter and stored in a sealed container. Animals frozen at the end of the metabolic testing were thawed and placed on pre-weighed (to 0.001 g accuracy) weigh boats prior to drying. Ochre stars were dried in a combustion oven (60°C) for 12 – 24 h until their dry mass had stabilized and then ashed in a muffle furnace (500°C) for 24 – 48 h. AFDM (*i.e.*, metabolically active tissue) was calculated as the dry mass (including the weigh boat) minus the ashed mass (including the weigh boat) while individual dry mass (*i.e.*, organic and inorganic mass) was calculated as the dry mass minus the weight of the empty weigh boat. The inorganic mass, primarily representing the ochre star’s calcified endoskeleton, was calculated as the ashed mass minus the weight of the empty weigh boat.

#### Data processing and statistical analyses

Absolute and mass-specific ṀO_2_ were calculated for each individual *Pisaster* using the “respR” package in R (Carey and Harianto, 2023). All ṀO_2_ measurements were adjusted for salinity and water volume in the chamber by correcting for each individual’s volumetric displacement. The ṀO_2_ values in the blank chamber of each run (< 0.1 mL O_2_/h) were used to correct for background respiration in the absolute and mass-specific ṀO_2_ values. Mass-specific values were adjusted for both wet mass and AFDM (*i.e.*, absolute ṀO_2_ divided by mass). The ṀO_2_ data was visually inspected to ensure that a linear decrease in water % air saturation of 5- 10% occurred and a minimum r^2^ of 0.98 was set to identify nonlinear measurements (Schuster *et al*., 2022). One measurement was discarded as it represented an extreme outlier in the data (n *=* 1/18; ṀO_2_ value 8x greater than all other values).

### Field observations

#### Natural moribundity in Pisaster

Field observations of moribund *Pisaster* (*i.e.*, individuals displaying loss of body turgor, visible lesions, and/or tissue degradation) were made at three sites in Barkley Sound (Eagle Bay, Grappler Narrows, and Strawberry Point; **Fig. 1**) in May, June and August 2023 for both juvenile and adult ochre stars to determine the natural prevalence of *Pisaster* moribundity in our study area. We surveyed ochre stars along walking belt transects from the waterline (at tidal datum of 0.33 m to 0.71 m) up to the highest occurring individual. *Pisaster* found within one meter of either side of the transect were counted and assessed for signs of moribundity: animals were recorded as “moribund” if white lesions were apparent on the body surface or if individuals displayed abnormal body turgor (Bates *et al*., 2009). A random sample of ∼25 ochre stars were measured for arm length along each transect, with 3-4 transects deployed at each site. Air temperature and SST was recorded at each site using a temperature probe (INKBIRD IBS-TH2 Plus) at the time of survey (**Table 1**).

#### Ethics statement

All research on *Pisaster* and *Mytilus* spp. was conducted under approval of the Department of Fisheries and Oceans Canada (Licence numbers XR 123 2023 and 134459), the Huu-ay-aht First Nations Department of Lands and Permitting (HFN permit number: 2023-017), and following the guidelines for animal care outlined by the Bamfield Marine Sciences Centre (BMSC) and the Canadian Council on Animal Care (CCAC).

## Results

### Experiment 1. Effects of the land-sea warming contrast on *Pisaster* feeding rate

#### Pisaster mortality

*Pisaster* mortality only occurred in the 30°C air treatments, with individuals in all other treatments surviving until the end of the experiment. Five (of 12) ochre stars in the 15°C water/30°C air treatment and three (of 12) ochre stars in the 20°C water/30°C air treatment died during the first half of the heat stress period (days 1 to 8); there was no further mortality in the 30°C air treatments after day seven of the heat stress period. *Pisaster* in the 30°C air treatments thus had a significantly higher probability of mortality (Cox proportional-hazards model; *p* = 0.01) than those in the 20°C or 25°C air treatments **(Fig. 2A)**. *Pisaster* mortality in the two 30°C air treatments did not differ, however, there was a trend towards higher mortality in the 15°C water treatment compared to the 20°C water treatment (**Fig 2A).** The probability of being consumed differed in mussels between treatments; mussels in the 15°C water/30°C air treatment experienced lower consumption than all other treatments (**Fig. 2B**; Cox proportional-hazards model; *p* = 0.04).

**Fig 2.**
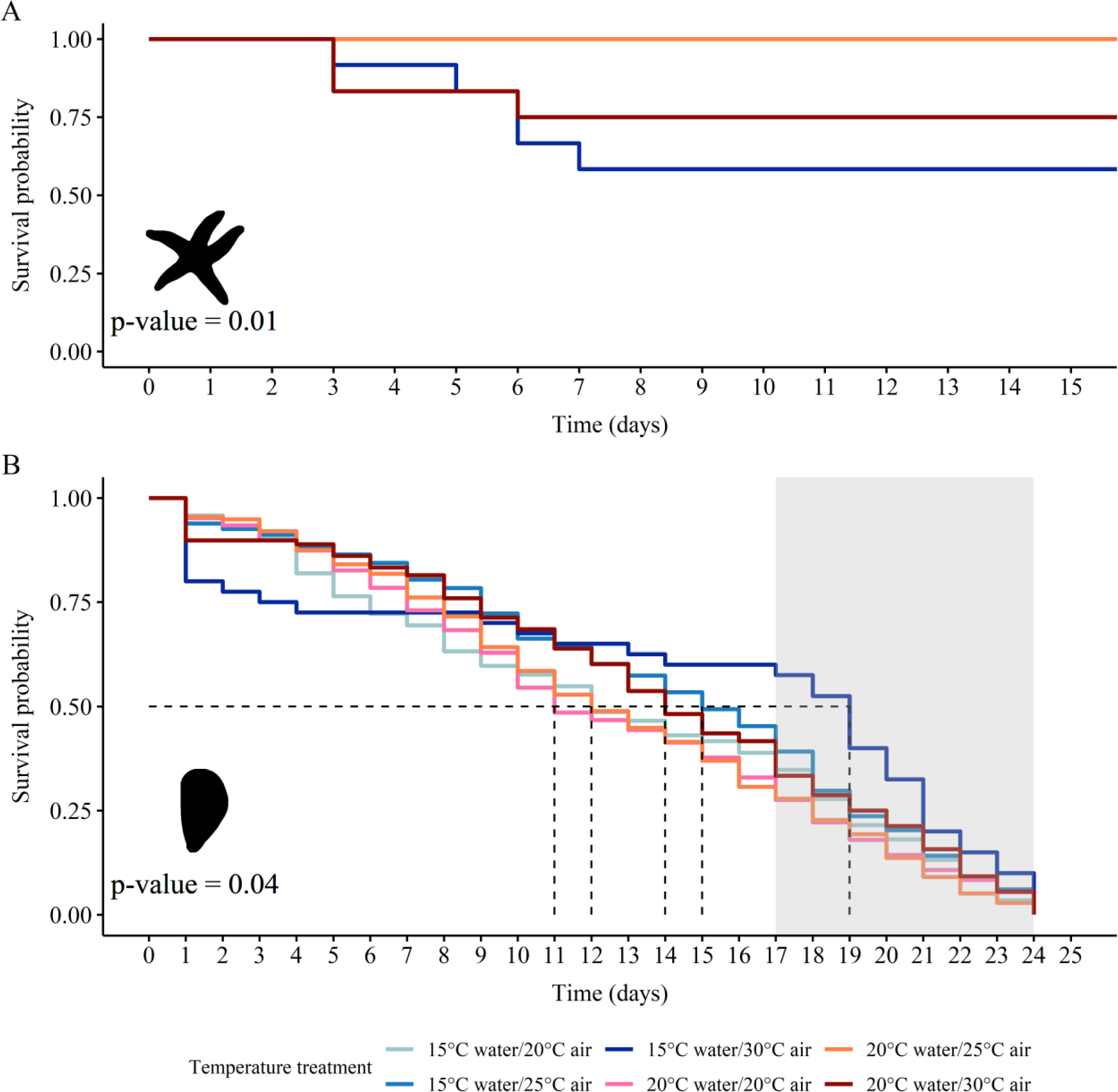
Kaplan-Meier survival curves for (A) juvenile *Pisaster ochraceus* (n = 72) and (B) consumed *Mytilus* spp. (n = 783) during the feeding activity experiment conducted in June 2023. White background shading represents the heat stress period of the experiment where sea stars were fed mussels *ad libidum* and alternated between water temperatures (15°C or 20°C) and air temperatures (20°C, 25°C, or 30°C) in a simulated high-tide (submerged for 18 hours) and low-tide (emersed for 6 hours) cycle. Grey background shading represents the recovery period of the experiment where sea stars were fully submerged in ambient water temperature (∼13°C) and continued to feed *ad libidum*. The mussels included in this analysis were all consumed by *Pisaster* during the experiment. **(A)** A significant difference in *Pisaster* survival probability was found between the 30°C air treatments compared with all other treatments (Cox proportional- hazards model; *p =* 0.01). Five *Pisaster* in the 15°C water/30°C air temperature treatment and three *Pisaster* in the 20°C water/30°C air temperature treatment died during the experiment. (B) *Mytilus* in the 15°C water/30°C air temperature treatment had a significantly higher survival probability than all other treatments (Cox proportional-hazards model; *p =* 0.04). The dotted lines represent the median survival probability for each temperature treatment.

#### Pisaster feeding activity

There was a significant difference in feeding activity (*i.e.*, the total number of mussels consumed by each juvenile *Pisaster*) between temperature treatments as determined by generalized linear mixed modelling (glmm). Overall, *Pisaster* in the 15°C water/30°C air treatment consumed fewer mussels than any other treatment across both the heat stress and recovery periods (**Table S3A, Fig. 3A**; *p* < 0.001). Additionally, these ochre stars consumed significantly more mussels during the recovery period than during the heat stress period (**Table S3A, Fig. 3A**; *p* = 0.001).

**Fig 3.**
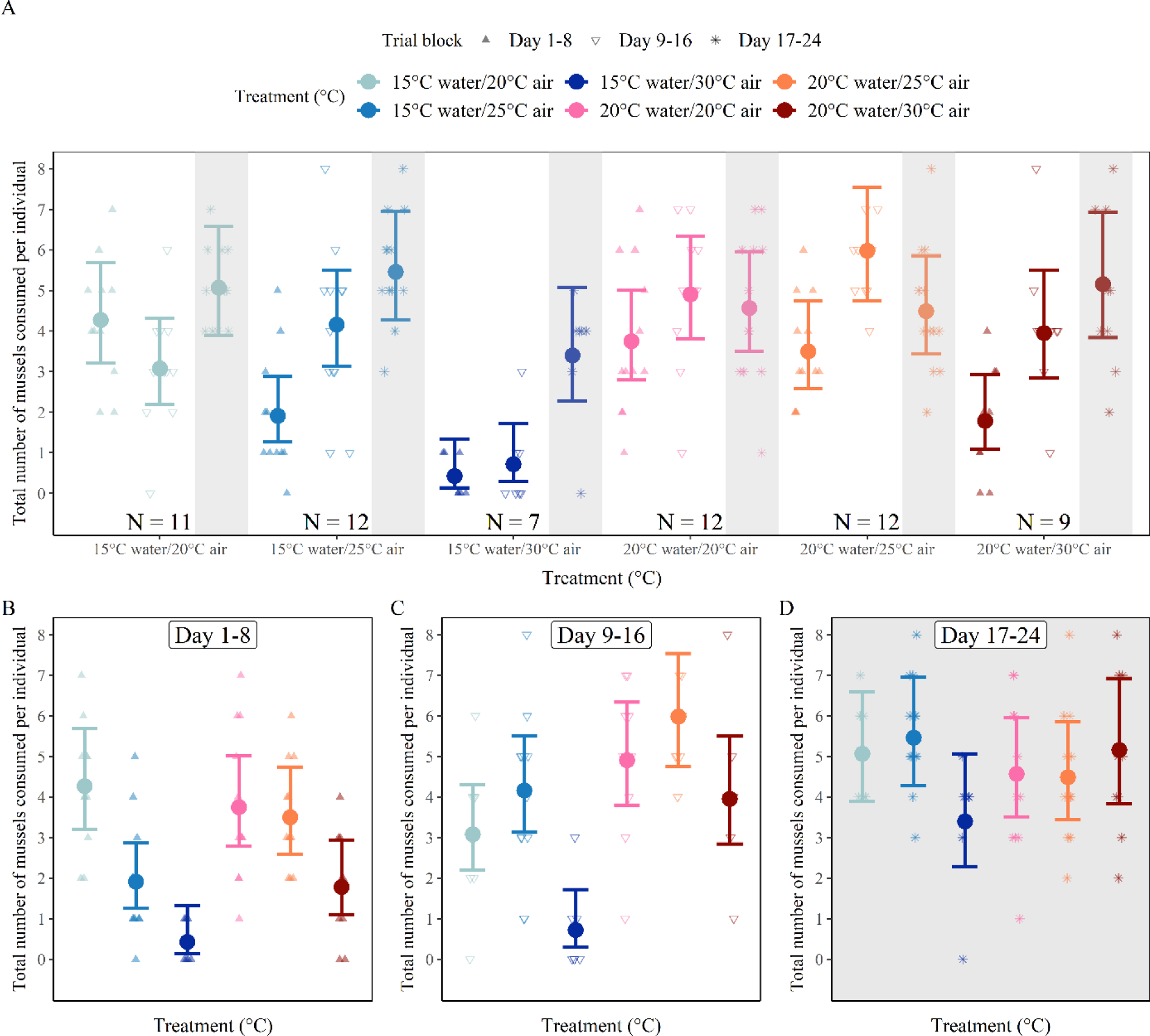
The total number of *Mytilus* spp. consumed by juvenile *Pisaster ochraceus* (n = 63) during the feeding activity experiment conducted in June 2023. White background shading represents the heat stress period of the experiment where sea stars were fed mussels *ad libidum* and alternated between water temperatures (15°C or 20°C) and air temperatures (20°C, 25°C, or 30°C) in a simulated high-tide (submerged for 18 hours) and low-tide (emersed for 6 hours) cycle. Grey background shading represents the recovery period of the experiment where sea stars were fully submerged in ambient water temperature (∼13°C) and continued to feed *ad libidum*. Dots and confidence intervals represent the model estimates produced by a generalized linear mixed model investigating differences in mussel consumption between treatments. Jittered points represent the total number of mussels consumed by each individual over three “blocks” of the experimental period. Each trial block was 8 days long, with days 1-8 representing the first half of the heat stress period (triangles), days 9-16 representing the second half of the heat stress period (open triangles), and days 17-24 representing the recovery period (asterisk). Sample sizes for each treatment are included in Panel A. Only sea stars that survived until the end of the experiment and retained five intact arms were included in analyses. **(A)** Total number of mussels consumed by *Pisaster,* grouped by treatment and separated by trial block. **(B, C, D)** Total number of mussels consumed by *Pisaster* in each trial block, grouped by treatment.

We also found differences in feeding activity within each 8-day experimental “time block”, *i.e.*, first half of the heat stress period: days 1-8; second half of the heat stress period: days 9-16; recovery period: days 17-24. During days 1-8 of the experiment, *Pisaster* in the 25°C and 30°C air treatments consumed fewer mussels than individuals in the 20°C air treatments (**Table S3B, Fig. 3B**; *p* < 0.01 and *p* < 0.001 respectively). A significant interaction was detected between water and air temperatures, with *Pisaster* in the 25°C and 30°C air treatments consuming more mussels in the 20°C water treatment than in the 15°C water treatment (**Table S3B, Fig. 3B**; *p* < 0.05). During days 9-16 of the experiment, *Pisaster* in the 20°C water treatments consumed more mussels overall than individuals in the 15°C water treatments, demonstrating a positive effect of water temperature on feeding activity (**Table S3C, Fig. 3C**; *p* < 0.05). Ochre stars in the 30°C air treatments still consumed significantly fewer mussels than individuals in the 20°C and 25°C air treatments, regardless of the water temperature (**Table S3C, Fig. 3D**; *p* < 0.01). There were no significant differences in feeding activity detected during the recovery period (days 17-24), however, there was a slight trend towards *Pisaster* in the 15°C water/30°C treatment consuming fewer mussels than all other treatments (**Table S3D; Fig. 3D)**.

Distinct shifts in *Pisaster* feeding activity (*i.e.*, mean number of mussels consumed per day) were detected using a change point analysis during the heat stress and recovery periods of the experiment. A change point was detected during the transition from the heat stress period to the recovery period (*i.e.*, days 16-18) for ochre stars in the 15°C water/30°C air treatment (**Fig. 4A**; changepoint detected on day 18). A one-way analysis of variance showed that ochre stars in this treatment (15°C water/30°C air) consumed significantly more mussels during the recovery period than in the first or second half of the heat stress period (**Fig. 4B**; Kruskal-Wallis test, *p* < 0.05). The change points for all other treatments were detected before the transition from the heat stress period to the recovery period and no further analyses were performed **(Fig. 4B)**.

**Fig 4.**
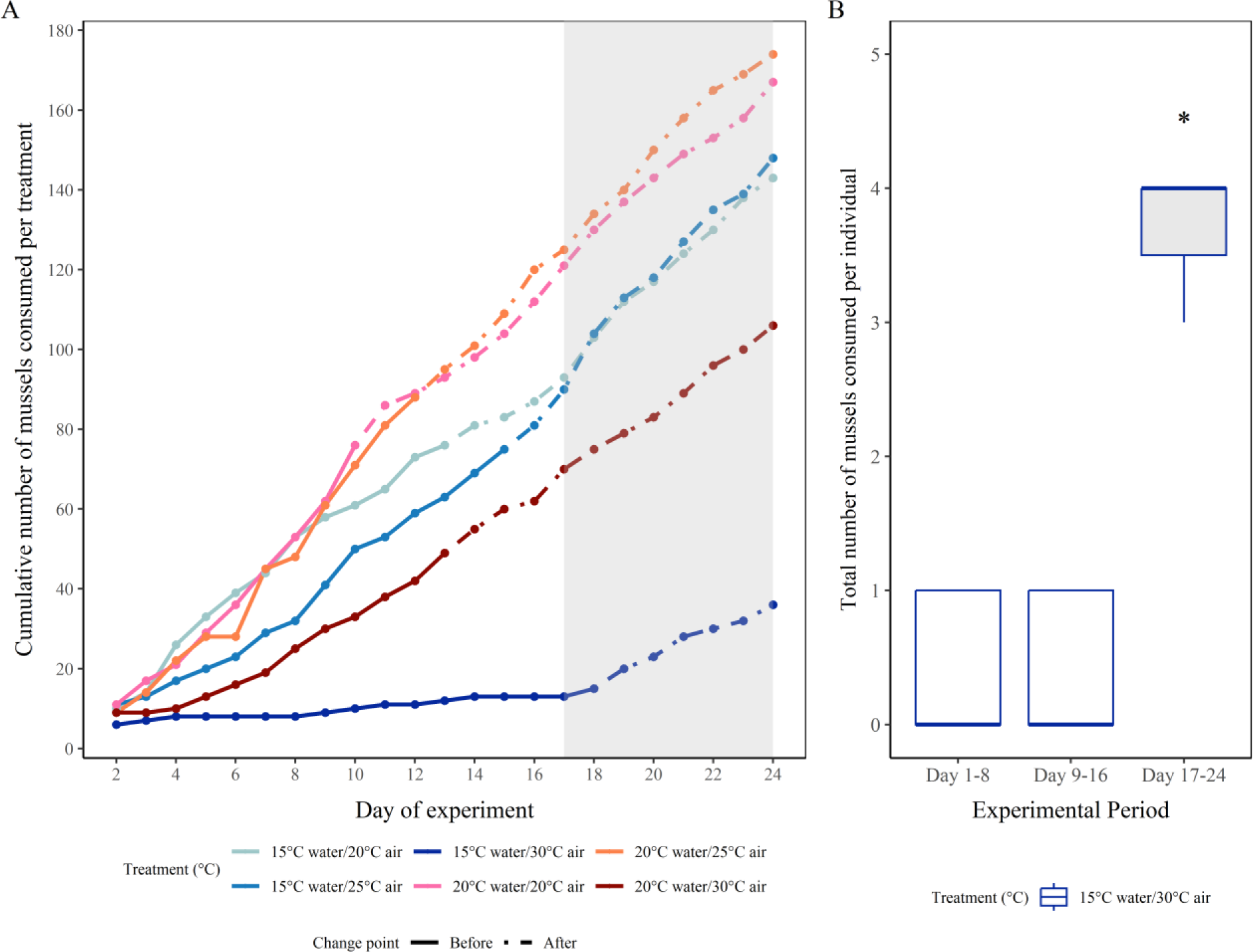
The cumulative number of *Mytilus* spp. consumed by juvenile *Pisaster ochraceus* (n = 63) during the feeding activity experiment conducted in June 2023. White background shading represents the heat stress period of the experiment where sea stars were fed mussels *ad libidum* and alternated between water temperatures (15°C or 20°C) and air temperatures (20°C, 25°C, or 30°C) in a simulated high-tide (submerged for 18 hours) and low-tide (emersed for 6 hours) cycle. Grey background shading represents the recovery period of the experiment where sea stars were fully submerged in ambient water temperature (∼13°C) and continued to feed *ad libidum*. **(A)** The cumulative number of *Mytilus* spp. consumed per treatment over the course of the experimental period. Change points (*i.e.*, changes in the mean number of mussels consumed) were detected for each treatment using the “changepoint” package in R. Solid lines represent consumption before the change point occurred while dot-dashed lines represent consumption after the change point was detected. **(B)** A change point was detected on day 18 (*i.e.*, during the shift from heat stress to recovery period) for *Pisaster* in the 15°C water/30°C air temperature treatment. The total consumption of mussels per *Pisaster* individual was grouped into three 8- day blocks representing the first half of the heat stress (days 1-8), the second half of the heat stress (days 9-16), and the recovery period (days 17-24). A significant difference in total mussel consumption was found between days 1-8 and days 17-24 (*p =* 0.02) and between days 9-16 and days 17-24 (*p =* 0.02).

### Experiment 2. Food availability, Pisaster metabolism, and air temperature

#### Pisaster mortality

No significant differences in *Pisaster* mortality were found between treatments, with all treatments exhibiting some level of mortality (**Fig. 5A**; Cox proportional-hazards model, *p* = 0.2). There was a trend towards higher mortality in fasted *Pisaster* compared to ochre stars that were fed *ad libidum* (**Fig. 5A**; +mussels/25°C = 6 individuals died, +mussels/30°C air = 11 individuals died, -mussels/25°C = 9 individuals died, -mussels/30°C = 12 individuals died).

**Fig 5.**
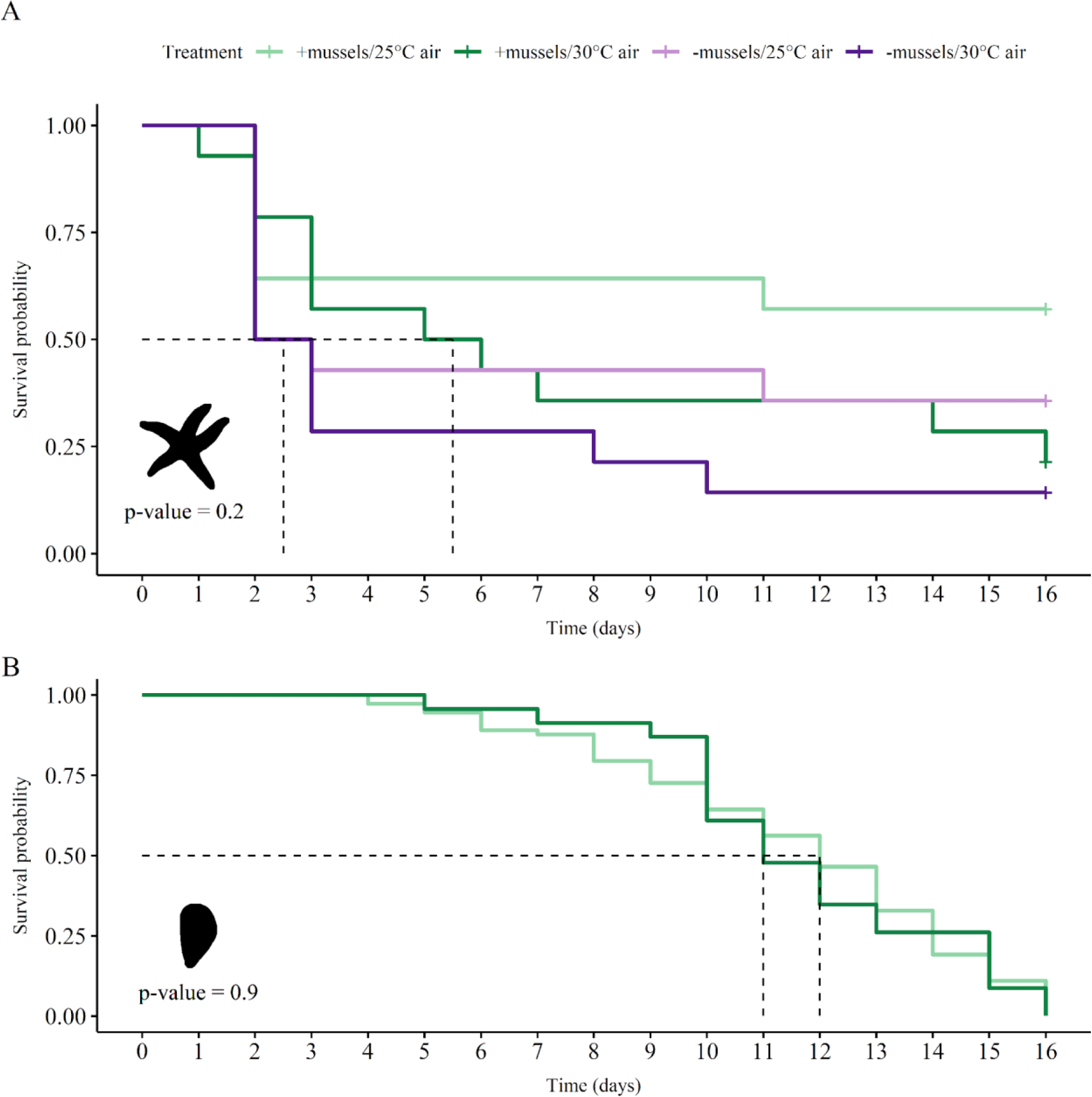
Kaplan-Meier survival curves for (A) juvenile *Pisaster ochraceus* (n = 56) and (B) consumed *Mytilus* spp. (n = 96) during the metabolic rate experiment conducted in August 2023. *Pisaster* were separated into four treatments (+mussels/25°C air, +mussels/30°C air, - mussels/25°C air, -mussels/30°C air). Individuals in the “+ mussels” treatments were fed mussels *ad libidum* for the duration of the experiment while individuals in the “- mussels” treatments were given no food (*i.e.*, fasted). *Pisaster* alternated between sea tables (20°C fixed water temperature) and incubators (25°C or 30°C air temperature) in a simulated high-tide (submerged for 18 hours) and low-tide (emersed for 6 hours) cycle. Dotted lines represent the median survival probability for each temperature treatment. **(A)** No significant difference in *Pisaster* survival probability was found between treatments (Cox proportional-hazards model; *p =* 0.2). A total of 38 individuals died over the course of the experiment (+mussels/25°C = 6 individuals died, +mussels/30°C air = 11 individuals died, -mussels/25°C = 9 individuals died, - mussels/30°C = 12 individuals died). **(B)** No significant difference in *Mytilus* survival probability was found between “+ mussels” treatments (Cox proportional-hazards model; *p =* 0.9).

#### Pisaster feeding activity

There was no significant difference in the probability of mussels being consumed between temperature treatments (**Fig. 5B**; Cox proportional hazards model, *p=* 0.9) and, analysing the data from a different perspective, no significant difference in *Pisaster* feeding activity (*i.e.*, the total number of mussels consumed by each juvenile ochre star) was found between treatments where mussels were provided *ad libitum* (**Fig. S1B;** t-test, *p* = 0.059). Even so, *Pisaster* in the +mussels/30°C air treatment did consume fewer mussels than those in the +mussels/25°C air treatment **(Fig. S1A, B)**. Lack of statistical power is expected as only 8 (of 14) individuals survived the +mussels/25°C air treatment and only 3 (of 14) individuals survived in the +mussels/30°C air treatment. Overall, *Pisaster* that consumed mussels during the experiment tended to have a lower probability of mortality compared with individuals that were fasted (**Fig. S2**).

#### Pisaster metabolic rate

In general, *Pisaster* did not differ in body mass or oxygen consumption between treatments. There was no difference found in wet mass (WM), ash-free dry mass (AFDM), or ADFM:WM ratio between treatments at any point of the experiment (ANOVA, *p* > 0.05). The ṀO_2_ per g of wet mass was significantly higher for *Pisaster* with access to mussels than for *Pisaster* that were fasted (Two-factor ANOVA, *p* = 0.02), with no differences found in ṀO_2_ between air treatments (**Fig. S3A)**. There were no significant differences detected in absolute ṀO_2_ or ṀO_2_ per g of AFDM between treatments (**Fig. S3B, C**).

### Comparison of mortality and feeding activity between experiments

Two treatments were repeated in both experiments: (*i.e.*, the 20°C water/25°C air/+mussels treatment and the 20°C water/30°C air/+mussels treatment). We thus compared the mortality and feeding activity of these treatment groups. Mortality was significantly higher during the second experiment compared with the first experiment **(Fig. 6A**; Cox proportional-hazards model, *p* < 0.001). Additionally, mortality was also higher in the 30°C versus the 25°C air treatments in experiments **(Fig. 6A**; Cox proportional-hazards model, *p* < 0.05). Feeding activity did not differ between the treatments or experiments **(Fig. 6B)**.

**Fig 6.**
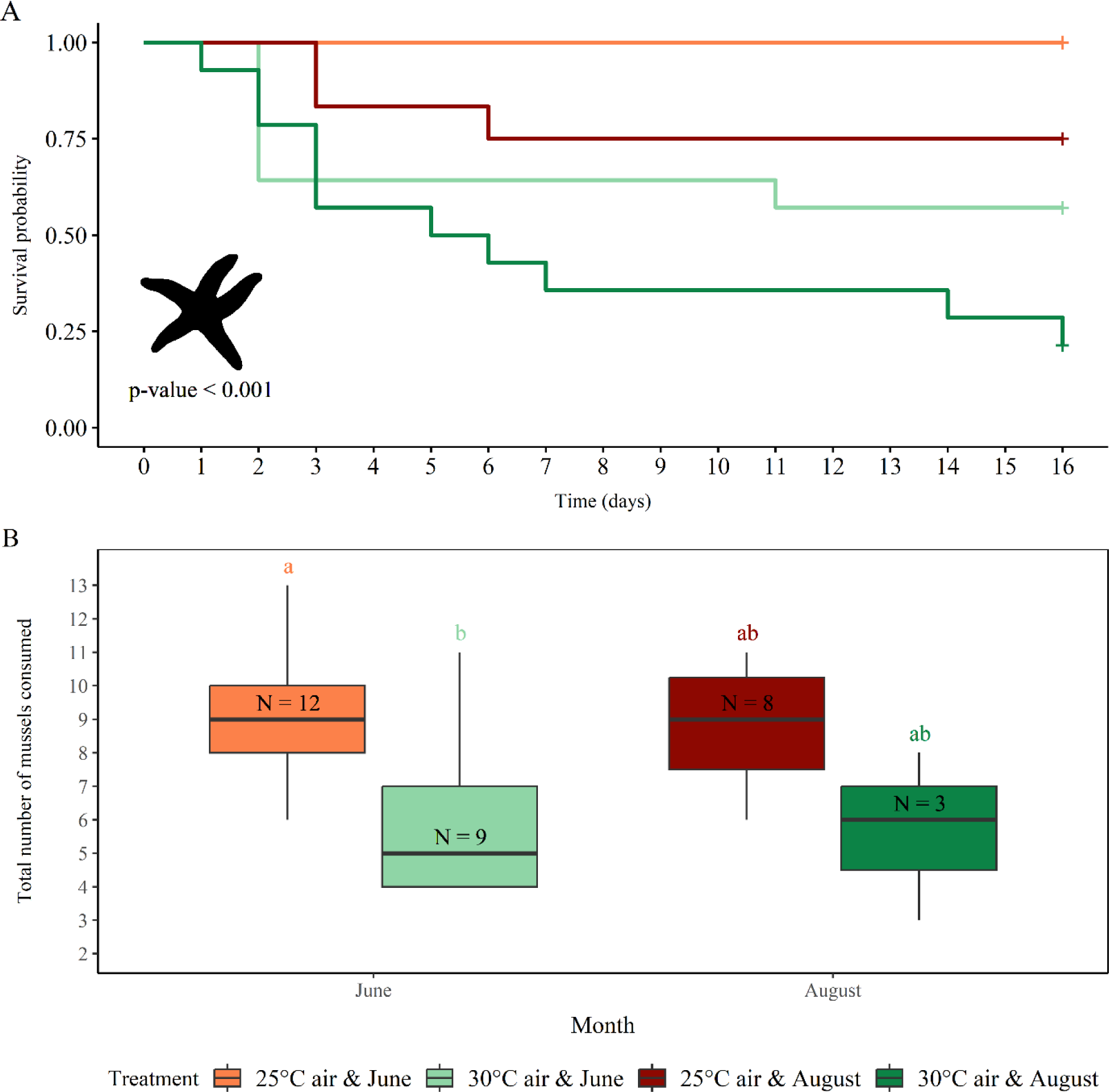
Comparison of juvenile *Pisaster ochraceus* survivorship (A) and feeding activity (B) between the feeding activity experiment conducted in June 2023 (Experiment 1) and the metabolic rate experiment conducted in August 2023 (Experiment 2). There were two treatments consistent across both experiments: 20°C water/25°C air/+mussels and 20°C water/30°C air treatments/+mussels. **(A)** Kaplan-Meier survival curve for *Pisaster* in June (n = 72) and August (n = 56). *Pisaster* in the second experiment (August) had a lower survival probability than those in the first experiment (June) (Cox proportional-hazards model, p < 0.001). *Pisaster* in the 30°C air treatments had a lower survival probability than individuals in the 25°C air treatments (Cox proportional-hazards model, p < 0.05). **(B)** The total number of mussels consumed by *Pisaster* in June (n = 21 individuals) and August (n = 11 individuals). There was no significant difference in feeding activity found between months, however, *Pisaster* in the 30°C air treatments tended to consumed fewer mussels than those in the 25°C air treatments.

### Field observations

#### Natural moribundity in Pisaster

All sites displayed very low levels of moribund *Pisaster* throughout the summer months, with the highest percentage of moribund individuals being 9.6% of adults in Eagle Bay in August **(Table 1)**. Overall, we observed the highest percentage of moribund adult and juvenile *Pisaster* at Eagle Bay, with signs of lesions and tissue degradation present in adults during all months and in juveniles during August. Moribundity was only recorded for adults and juveniles at Grappler Narrows in August, and no moribund individuals were present at Strawberry Point for any month surveyed (**Table 1**). Air temperatures ranged from 15.4 °C (May) to 21.8 °C (June) at Eagle Bay, 17.8 °C (May) to 23.5 °C (August) at Grappler Narrows, and 14.3 °C (June) to 21 °C (May) at Strawberry Point while SST ranged from 14.7 °C (May) to 19.5 °C (August) at Eagle Bay, 14.6 °C (June) to 17.6 °C (August) at Grappler Narrows, and 14.7 °C (June) to 17.5 °C (August) at Strawberry Point (**Table 1**).

## Discussion

Here we find that cooler ocean temperatures in the spring and summer may cause physiological stress for juvenile *Pisaster* exposed to warm air. Indeed, animals exposed to both the coolest seawater (∼15°C) and warmest air (∼30°C) temperatures in our study exhibited the highest levels of mortality and consumed the least number of mussels overall. We further found a positive effect of the 20°C ocean temperatures on *Pisaster* feeding rate regardless of air temperature. Our results thus show that changes in water and air temperature have a strong influence on juvenile *Pisaster* feeding rates and mortality. We further find, in this particular case, rather than creating thermal refugia, relatively cooler ocean temperatures compared to air broadens the range of temperatures experienced by species using both habitats, which may lead to decreases in performance. Our findings indicate that intertidal organisms which adjust thermal tolerance limits to accommodate cooler ocean temperatures when submerged may be disadvantaged if extreme heat stress occurs during aerial emersion.

Our experiments indicate juvenile *Pisaster* are sensitive to thermal stress. Elevated aerial temperatures (∼30°C) typical of extreme heat events increased mortality and decreased the feeding rate of juvenile ochre stars. These results presumably indicate juvenile *Pisaster* in the ∼30°C air treatments surpassed their thermal optimum (*T_opt_*) and were nearing their critical upper thermal limit (Pincebourde *et al*., 2008; Brown *et al*., 2009). The aerial lethal limit (*i.e.*, quantified here as the LT50; the point at which 50% of individuals die following heat exposure) of adult *Pisaster* is close to 35°C, with virtually no mortality occurring at body temperatures below this threshold when tested experimentally (Pincebourde *et al*., 2008). Mortality in juvenile *Pisaster* began around 30°C which reveals a greater sensitivity to thermal stress for this early life stage (assuming that populations have similar thresholds in different years). In fact, a similar ontogenetic shift in stress tolerance has been observed in other species of intertidal invertebrates (*e.g.*, mussels, barnacles, snails; Jenewein and Gosselin, 2013; Hamilton and Gosselin, 2020) with organisms generally becoming more tolerant to elevated temperatures and desiccation as they move from juvenile to adult stages. Differences in thermal tolerance between life stages may relate to microhabitat use, as several species of intertidal animals live in more protected areas (*i.e.*, filamentous algae, mussel beds, underneath rocks) during early benthic phases before moving to more exposed microhabitats as late-stage juveniles or adults (Hunt and Scheibling, 1996; Gosselin, 1997; Jenewein and Gosselin, 2013). *Pisaster* likely rely on behavioural adaptations, such as seeking out sun-protected microhabitats, to mitigate the negative physiological effects of high temperatures during emersion (Petes *et al*., 2008; Szathmary *et al*., 2009). Indeed, most juveniles in the present study were observed underneath rocks or deep in crevices where sun exposure was minimal. The presence of suitable microhabitats will thus be critical for *Pisaster* persistence under climate warming, especially for the more thermally susceptible juvenile life stage.

The thermal sensitivity of juvenile *Pisaster* was further evidenced by complex feeding responses during prolonged aerial heat stress. First, mussel consumption increased from the ∼20°C to ∼25°C air treatments, but declined significantly at ∼30°C. The higher feeding observed in the ∼25°C air treatments compared to the ∼20°C air treatments may reflect a greater need for prey assimilation to compensate for the energetic costs of a higher metabolism under warming conditions (Pincebourde *et al*., 2008; Fly *et al*., 2012). Moreover, in the ∼30°C air treatments juvenile *Pisaster* had significantly reduced feeding rates, which may indicate a decrease in metabolic activity as individuals surpass their thermal optimum (*T_opt_*) and move closer to their critical upper limit (*CT_max_*), a response that has also been observed in adult ochre stars exposed to chronic aerial thermal stress (Pincebourde *et al*., 2008).

The duration of aerial heat stress also influenced feeding in juvenile *Pisaster,* with greater rates of mussel consumption in the latter portion of the heat stress period. Rapid shifts in mussel consumption (*i.e.*, change in the mean number of mussels consumed, evidenced by a change point analysis) were observed past the midway point (*i.e.*, day 8 of the experiment) of the heat stress period (Experiment 1) in all treatments. *Pisaster* likely increased their feeding rates as the heat stress period progressed to compensate for the prolonged physiological stress and elevated metabolic demands at higher temperatures. Contrary to our expectations, ochre stars exposed to the coolest water (∼15°C) and warmest air (∼30°C) did not shift their feeding rate until the heat stress had ended, and continued to consume fewer mussels than all other treatments during the recovery period. Thus, prolonged aerial thermal stress combined with cooler water temperatures led to negative performance in juvenile *Pisaster*.

Warming water temperatures positively affected feeding rates in juvenile *Pisaster,* and may help compensate for the energetic demands of aerial heat stress. Overall, the treatment with the greatest temperature contrast (*i.e.*, 15°C water/30°C air treatment) led to the lowest number of mussels consumed across all treatments and the highest mortality. More mussels were consumed by juvenile *Pisaster* experiencing ∼20°C water temperatures compared to those in 15°C water temperatures and higher mortality also occurred in the 15°C water temperatures compared to the warmer water (∼20°C) temperatures for individuals in the ∼30°C air treatments. Warming water temperatures, like air temperatures, elevate metabolic activity and may prompt increased feeding rates to compensate for associated energetic costs (Pincebourde *et al*., 2008; Fly *et al*., 2012). Juvenile *Pisaster* in the 20°C water/25°C air treatment consumed the most mussels overall, indicating that warm water and warm air can have a positive effect on feeding rate when temperatures fall within an optimal thermal range. Warmer ocean conditions may also ameliorate the negative impacts of aerial heatwaves, as juveniles in warmer water (∼20°C) had significantly higher feeding rates than individuals in cooler water (∼15°C) when exposed to 30°C air temperatures. Thus, juvenile ochre stars may be particularly vulnerable to the “land-sea warming contrast”, as faster warming on land leads to hotter aerial temperatures in the spring when ocean conditions are still relatively cooler.

We further found that experimental mortality (*i.e.*, deaths recorded during our experiments) and natural moribundity (*i.e.*, percent of moribund individuals observed in the field) were highest in late summer. Juvenile *Pisaster* mortality was significantly higher in Experiment 2 (August; 61% mortality, n = 34/56) with moribund individuals recorded in all treatments, compared to Experiment 1 (June; 13% mortality, n = 8/72) where only individuals subjected to the warmest air (∼30°C) died. One explanation is that organisms may accumulate energetic debt in combination with physiological stress damage which can compromise survival and physiological performance after prolonged summer conditions (Dahlhoff *et al*., 2001). The mortality observed in our August experiment was mirrored by juvenile *Pisaster* moribundity at Eagle Bay in August, although at much lower rates (8.3% moribund juveniles in August versus 0% earlier in the year). The lower rates of juvenile moribundity in the field may be affected by detection biases due to their small size and potential to be washed away during high tides (Menge *et al*., 2016). Additionally, temperatures in the field may not have reached the extreme aerial conditions (∼30°C) present in the experiments which led to the highest rates of juvenile mortality.

There are several implications of *Pisaster* feeding rates changing in response to different temperature combinations during ocean and air exposure. Our cooler water temperature (∼15°C) treatments were consistent with sea surface temperatures recorded near Bamfield in May and June, while our warmer water temperature (∼20°C) treatments generally represented sea surface temperatures in August. Average temperatures of the sea surface and land surface have increased over time, albeit at differing rates according to the “land-sea warming contrast” (Sutton *et al*., 2007; Joshi *et al*., 2013). As average ocean and air temperatures increase, fluctuations in temperature become greater and extreme heat events more likely (IPCC, 2023). Thus, extreme heat events in the spring and summer may increase in frequency and drive higher aerial exposure that is damaging to organisms exposed during low tide. In fact, our results suggest that early summer (May and June) heatwaves, with cooler ocean temperatures and high air temperatures, will have the greatest negative impact on survival and feeding activity in juvenile *Pisaster*. This is shown directly in our 15°C water/30°C air treatment which had significantly higher levels of mortality and lower levels of mussel consumption than our other experimental treatments.

Juvenile *Pisaster* may adjust their upper tolerance limits to accommodate cooler ocean temperatures in early summer, and the physiological trade-off may be greater sensitivity to high aerial temperatures when emersed. We also suggest that juvenile *Pisaster* will be in poorer physiological condition during the late summer months due to physiological stress damage accumulated throughout the summer. This is evidenced by the lasting negative impacts of the 15°C water/30°C air treatment on feeding in our first experiment and the increased mortality rates recorded during our second experiment.

Reduced feeding throughout the summer may have lasting effects on *Pisaster* population structure due to seasonal reproduction. Adult *Pisaster* spawn in spring, coinciding with phytoplankton blooms that support larval development, and settlement happens approximately two months later, giving juvenile *Pisaster* the opportunity to feed all summer and allocate any surplus of energy towards growth (Mauzey, 1966; Robles, 2013; Robles *et al*., 2021). Once the size for sexual maturity is reached (*i.e.*, between 70 and 150g; Robles, 2013), the energy assimilated from active foraging is stored in the pyloric caeca and used for gonad development in the winter. The mass of the pyloric caeca prior to winter dormancy is almost proportional to the mass of gametes released during the next spring spawn, meaning *Pisaster* reproductive output is largely determined by summer feeding rates (Sanford and Menge, 2007; Robles *et al*., 2021). If an aerial heatwave hits ochre star populations in early summer when water temperatures remain cooler, subsequent reductions in feeding rates may slow the growth rate of juveniles and delay the onset of reproductive maturity. Increased rates of juvenile mortality under these same conditions may limit population growth as fewer individuals survive to the adult life stage. For adult ochre stars, reduced feeding due to thermally stressful conditions in the summer may also decrease reproductive output the following year, leading to lowered levels of recruitment and settlement. Thus, the “land-sea warming contrast” may interact with phenological patterns and has the potential to severely diminish the reproductive potential of *Pisaster* populations as well as overall population size through its effects on physiology and feeding behaviour.

In addition to population-level impacts, greater differences in temperature between habitats that cross the ocean-air interface may cause community-level changes through alterations in keystone predation (*i.e.*, species that have a disproportionate effect on a community relative to its biomass; Paine, 1966). When ocean and air temperatures warm, but remain within the optimal thermal range of juvenile *Pisaster*, feeding increases to compensate for the energetic demands of a higher metabolic rate. Thus, a larger proportion of mussels, and their associated biodiversity (*i.e.*, infauna, epifauna, pelagic fauna), may be consumed by ochre stars during minor heat events or seasonal warming of the air (∼20 to 25°C). In this case, keystone predation may shift community structure by clearing mussel beds, which provide critical habitat space for many organisms (Beadman *et al*., 2004; Benjamin *et al*., 2022). Mussel populations are also sensitive to elevated aerial temperatures, and lethal levels of desiccation stress for recently settled *Mytilus* spp. can occur regularly during recruitment season in Barkley Sound (Jenewein and Gosselin, 2013). Therefore, increased temperature may have a two-fold effect on the *Pisaster*-*Mytilus* trophic interaction as greater predation rates occur simultaneously with a low recruitment of mussels, generating a positive feedback loop that destabilizes both predator and prey populations. If ocean and air temperatures surpass the upper critical limits of juvenile *Pisaster*, feeding rates may decrease to free up oxygen for processes other than digestion. Thus, under relatively cool ocean temperatures and elevated air temperatures, small mussels may be released from predation pressure, thereby reducing intertidal species diversity as they outcompete other sessile organisms for space. Such shifts in benthic community composition have already been observed through the removal of adult *Pisaster* (Paine, 1966, 1969; Robles, 2013; Robles *et al*., 2021), however, community responses to changes in juvenile ochre star behaviour have yet to be properly investigated.

## Conclusion

Physiological and behavioural responses of a predatory sea star to experimentally manipulated water and air temperatures implicate the role of increased temperature contrast between the land and sea in impacting the feeding behaviour and survivorship of this keystone species. This highlights an important gap in our understanding of how other taxa may respond to differing rates of land and sea warming, particularly when cooler ocean temperatures occur simultaneously with extreme aerial heat events. Additionally, comparisons of physiological and behavioural responses to thermal stress between life stages will be key in predicting population-level impacts as juveniles may be more thermally sensitive than adults. Understanding the impacts of different patterns of ocean and aerial temperature exposure will underpin shifts in species interactions and community dynamics.

## Acknowledgements

We thank the staff and researchers at the Bamfield Marine Sciences Centre (BMSC) for hosting our research and providing essential resources for our experiments. We thank Mike Delsey for building the temperature-controlled incubators used in our experiments. We also thank Mara Bohm, Emily Braun, Valesca de Groot, and Logan O’Reilly for assisting with animal collections and field surveys. We thank Julia Baum, Matthew Csordas, Chris Neufeld and Sam Starko for contributing the air temperature data from Eagle Bay.

## Data availability

All relevant data available on GitHub (https://github.com/lydiawalton).

## List of Symbols and Abbreviations

*CTmin*: critical lower thermal limit of an organisms’ thermal range
*Topt*: thermal optimum of an organisms’ thermal range
*CTmax*: critical upper thermal limit of an organisms’ thermal range
*ṀO2*: oxygen consumption of an organism (metabolic rate)
*AFDM*: ash-free dry mass (the amount of organic, or metabolically active tissue, in an organism)

